# Major complex trait for early de novo programming ‘CoV-MAC-TED’ detected in human nasal epithelial cells infected by two SARS-CoV-2 variants is promising for designing therapeutic strategies

**DOI:** 10.1101/2021.10.19.464952

**Authors:** José Hélio Costa, Shahid Aziz, Carlos Noceda, Birgit Arnholdt-Schmitt

## Abstract

**Background:** Early metabolic reorganization was only recently recognized as essentially integrated part of immunology. In this context, unbalanced ROS/RNS levels that connected to increased aerobic fermentation, which linked to alpha-tubulin-based cell restructuration and control of cell cycle progression, was identified as *ma*jor *c*omplex *t*rait for *e*arly *d*e novo programming (‘CoV-MAC-TED’) during SARS-CoV-2 infection. This trait was highlighted as critical target for developing early anti-viral/anti-SARS-CoV-2 strategies. To obtain this result, analyses had been performed on transcriptome data from diverse experimental cell systems. A call was released for wide data collection of the defined set of genes for transcriptome analyses, named ‘ReprogVirus’, which should be based on strictly standardized protocols and data entry from diverse virus types and variants into the ‘ReprogVirus Platform’. This platform is currently under development. However, so far an *in vitro* cell system from primary target cells for virus attacks that could ideally serve for standardizing data collection of early SARS-CoV-2 infection responses was not defined.

**Results:** Here, we demonstrate transcriptome level profiles of the most critical ‘ReprogVirus’ gene sets for identifying ‘CoV-MAC-TED’ in cultured human nasal epithelial cells. Our results (a) validate ‘Cov-MAC-TED’ as crucial trait for early SARS-CoV-2 reprogramming for both tested virus variants and (b) demonstrate its relevance in cultured human nasal epithelial cells.

**Conclusion:** *In vitro*-cultured human nasal epithelial cells proved to be appropriate for standardized transcriptome data collection in the ‘ReprogVirus Platform’. Thus, this cell system is highly promising to advance integrative data analyses by help of Artificial Intelligence methodologies for designing anti-SARS-CoV-2 strategies.

## 1. Background

### Is there a paradigm shift in understanding immunology?

It is increasingly understood that plants and animals have similar responses and cell memory mechanisms to manage immunology (1, 2). Effective immunologic protection requires a diversity of innate and adaptive cell responses and cell memory tools (1–4). However, immunologic responses are energy consuming and require efficient metabolic reprogramming, but metabolic reorganization is not fully recognized as an integrated part of immunology (5–8). Viruses have comparatively low Gibbs energy due to their chemical compositions (9, 10). This makes their replication highly competitive and is the basis for their ‘structural violence’ against the host cell metabolism [see discussion on this term in Costa et al., 2021 (11)]. Virus-induced host cell reprogramming that is intended to support host defense is turned ad absurdum for the host and favors virus replication as the driving force. As a consequence of this conflict, any viral infection provokes struggling for commanding coordination of host cell program and this starts in the initially infected cells. Competing for bioenergy and for ‘territories’ is decisive for the success of virus reproduction and evolution.

These insights stimulated our group to take profit of mixed plant and human cell-based experimental systems for gaining general, relevant knowledge on early reprogramming during SARS-CoV-2 infection. Our complex approach was explained on the basis of extensive literature reviews in a recent paper entitled ‘From plant survival under severe stress to anti-viral human defense – a perspective that calls for common efforts’ (11, 12). Having applied our theoretical concept by comparing transcriptome data from a resilient plant system with coronavirus-infected human cells of diverse origins, a *ma*jor *c*omplex *t*rait for *e*arly *d*e novo programming upon SARS-CoV-2, named ‘CoV-MAC-TED’, was identified that should help tracing critical virus footprints. This trait covers unbalanced levels of ROS/RNS (i.e. reactive oxygen species in relation to reactive nitrogen species), which connects to temporarily increased aerobic fermentation that links to α-tubulin-based cell restructuration and cell cycle regulation (11).

### Resilience can depend on the capacity for efficient early reprogramming – learning from plants

Plants as settled organisms are especially challenged to rapidly confront highly diverse and complex abiotic and biotic environmental constraints, including virus threats. Marker development is essentially required to permanently advance breeding strategies that can timely cope with ever changing environmental conditions, such as climate changes. Consequently, prediction of adaptive robustness that provides resilience is best explored in plants (13). In this context, early reprogramming in target cells and tissues that impacts final agronomic or quality characteristics (such as yield stability or richness in secondary metabolites) was raised by our group as a trait *per se* (14–20). We developed unique concepts and tool kits that allow predicting plant robustness in the field from respiration traits as early as several hours during seed germination (21–26). This innovative approach was tested and preliminary validated by using diverse plant species (13, 25–26). This observation suggests common mechanisms for resilient life performance across plant species. As a hypothesis, we expect similar performance across oxygen-dependent, respiring eukaryotic organisms. Applying a higher degree of abstraction, we argued that these results might also be promising to identify critical mechanisms in human cells under virus stress including SARS-CoV-2, which might help to pave the way for designing early combating strategies (11, 12).

### Early reprogramming can link to ROS/RNS equilibration and sugar-dependent fermentation

We found that adaptive early reprogramming in plants can be critically connected to temporarily enhanced, sugar-dependent fermentation (11, 26). This process was essentially regulated by an enzyme named alternative oxidase (AOX), which maintains ROS and RNS levels equilibrated, regulates metabolic and energetic homeostasis and adjusts respiration overload and fermentation, which at the same time associates to induction and regulation of cell division growth (11, 26). These characteristics make AOX a highly promising functional marker resource for improving general plant resilience. Its role in early reprogramming was indicated in several applied systems cited in Arnholdt-Schmitt et al. (12). However, humans do not possess this enzyme (see discussions in 11, 12). Nevertheless, understanding the functional importance of AOX in plants can guide research strategies to identify marker gene candidates for resilience in human target cells against virus attacks appropriate to characterize resilient traits early at reprogramming. Costa et al. (11) suggested that melatonin could at least in part substitute the early role of AOX during reprogramming. Melatonin is a natural hormone in humans, which has also been recognized as a phytohormone (27). It is produced in most organs and cells (27–31), including human salivary gland cells (32). Costa et al. (11) observed that *ASMT* transcript levels were increasing dependent on MOI level and infection time in MERS-CoV-infected MRC5 cells, which encouraged studying markers for melatonin metabolism in nose cells. Melatonin is known to possess anti-oxidant and pathogen defense-related properties and shows high fluctuation in its cellular concentration (27, 33–37). Furthermore, melatonin is since long proposed as anti-viral agent (38) promising as repurposing drug to treat SARS-CoV-2 infections (39, 40).

### Driving a standardized collection of data on virus-induced early reprogramming

Arnholdt-Schmitt et al. (12) initiated common efforts for a standardized collection of transcriptome data during early reprogramming after virus infection from a defined set of genes, named ‘ReprogVirus’. The principle intention of this approach was to identify common early target traits that could help in designing therapeutic strategies, which could be applied for a high diversity of virus types and variants. In the present communication, we reduced the number of tested genes from ReprogVirus to promising core markers for CoV-MAC-TED components (11). Thus, genes were selected to identify a shift in ROS/RNS (ASMTL, SOD1, SOD2, ADH5, NOS2), represent glycolysis (PFK, GAPDH, Eno) and lactic acid fermentation (LDH) as well as structural cell organization (α-Tub). We assumed that ASMTL could indicate oxidative stress equilibration, while SOD1 and SOD2 genes mark anti-oxidative activities and were selected to indicate oxidative stress. ASMTL is a paralog of ASMT, which is involved in melatonin synthesis in human cells (28). However, we could not find ASMT gene transcripts in collected human nasal epithelial cells. ADH5 is known to be involved in ROS/RNS equilibration through NO homeostasis regulation (40–42) and the inducible NOS gene, NOS2, relates to the induction of NO production (11, 43). NOS1 and NOS3 had not been encountered in sufficient quantities in the collected epithelial nose cell data. Further, we selected SNRK and mTOR to highlight cell energy-status signaling (44–47). The mTOR is activated when there is excess of energy in contrast to SNRK, where higher expression indicates energy depletion. Genes for E2F1 together with mTOR were included to indicate changes in cell cycle regulation, namely cell cycle progression (G1/S and G2/M transitions) (47). E2F1 belongs to the transcription factor family E2F and is known as cell cycle activator. The interferon regulator factor, IRF9, demonstrated early transcription in SARS-CoV-2 infected human lung adenocarcinoma cells and was therefore proposed as functional marker candidate that could signal initiation of the classical immune system (11). Here, we studied validity of our approach for SARS-CoV-2 in human nasal epithelial cells (NECs) that was originally identified to cause the Coronavirus Disease 2019 (COVID-19). Additionally, we tested whether the same approach could be applied to a SARS-CoV-2 variant (SARS-CoV-2 Δ382). This mutant was detected in Singapore and other countries and had been associated with less severe infection (48, 49).

## 2. Material and Methods

### 2.1 Gene expression analyses of RNA-seq data from SARS-CoV2 infected human nasal epithelial cells

In this work we analyzed the expression of the main ReprogVirus genes (11, 12, for gene abbreviations see in supplementary table S1) in transcriptomic data from human nasal epithelial cells infected with two SARS-CoV2 variants (wild type and mutant Δ382) at 0, 8, 24 and 72 hpi (hours post infection). Transcriptomic data (RNA-seq data) are available in SRA database at GenBank (NCBI) under the bioproject PRJNA680711 previously published by Gamage et al. (48). The ReprogVirus genes analyzed were involved in shift in ROS/RNS (ASMTL, SOD1, SOD2, ADH5, NOS2), glycolysis [Total PFK (PFKM, PFKL and PFKP), GAPDH, Total Eno (Eno1, Eno2 and Eno3)], lactic acid fermentation [Total LDH (LDH-A, LDH-B, LDH-C, LDH-AL6A, LDH-AL6B)] structural cell organization [Total α-Tub (TUB-A1B, TUB-A1C, TUB-A4A)], cell cycle activator (E2F1), cell energy-status (SNRK, mTOR) and immune response (IRF9). Accession numbers of these genes are available in Costa et al. (11). SARS-CoV-2 proliferation was monitored evaluating transcript levels of virus helicase (YP_009725308.1) and virus RNA-dependent RNA-polymerase (RdRp) (YP_009725307.1) genes.

Gene expression was evaluated by mapping, quantifying and normalizing the reads of each ReprogVirus gene in the RNA-seq data with three biological replicates. For this, ReprogVirus cDNAs were aligned against RNA-seq data using the Magic-Blast software (50). Specific parameters in Magic-Blast as word size of 64 was included to certificate specific read detection for each gene. The number of mapped reads was obtained using the HTSeq (51) program exploring an alignment file (in SAM format) derived from Magic-Blast. Normalization of reads was performed using the RPKM (Reads Per Kilobase of transcript per Million of mapped reads) method (52) according to the following equation: RPKM = (number of mapped reads X 10^9^) / (number of sequences in each database X number of nucleotides of each gene).

### 2.2. Statistical analyses

Normality and homogeneity of variances from the analyzed variables (in RPKM) were tested with Shapiro-Wilk test and Levene tests, respectively, using InfoStat 2018I. Then, ANOVA tests for single measures were performed for each donor and virus variant along different time points by using Excel datasheet. Significance levels were set at α=0.05.

We highlight that we interpret our data as ‘real’ observations under the employed conditions involving only small samples, which certainly provide insights that cannot get relevance or not relevance by using significance calculation. Nevertheless, we applied significance calculations at usual p-values for biological research as an aid to focus our insights. Readers are encouraged to making themselves familiar with the current paradigm change related to the usage of statistical significance (53–57).

## 3. Results

**Figure 1** shows relative transcript level changes of selected ReprogVirus genes in epithelial nasal cells infected with SARS-CoV-2 or SARS-CoV-2 mutant Δ382 at 8 hpi, 24 hpi and 72 hpi. Transcript level changes are expressed as % of 0 hpi. Standard errors (SE) are listed for all genes in **supplementary table S1**.

**Figure 1:**
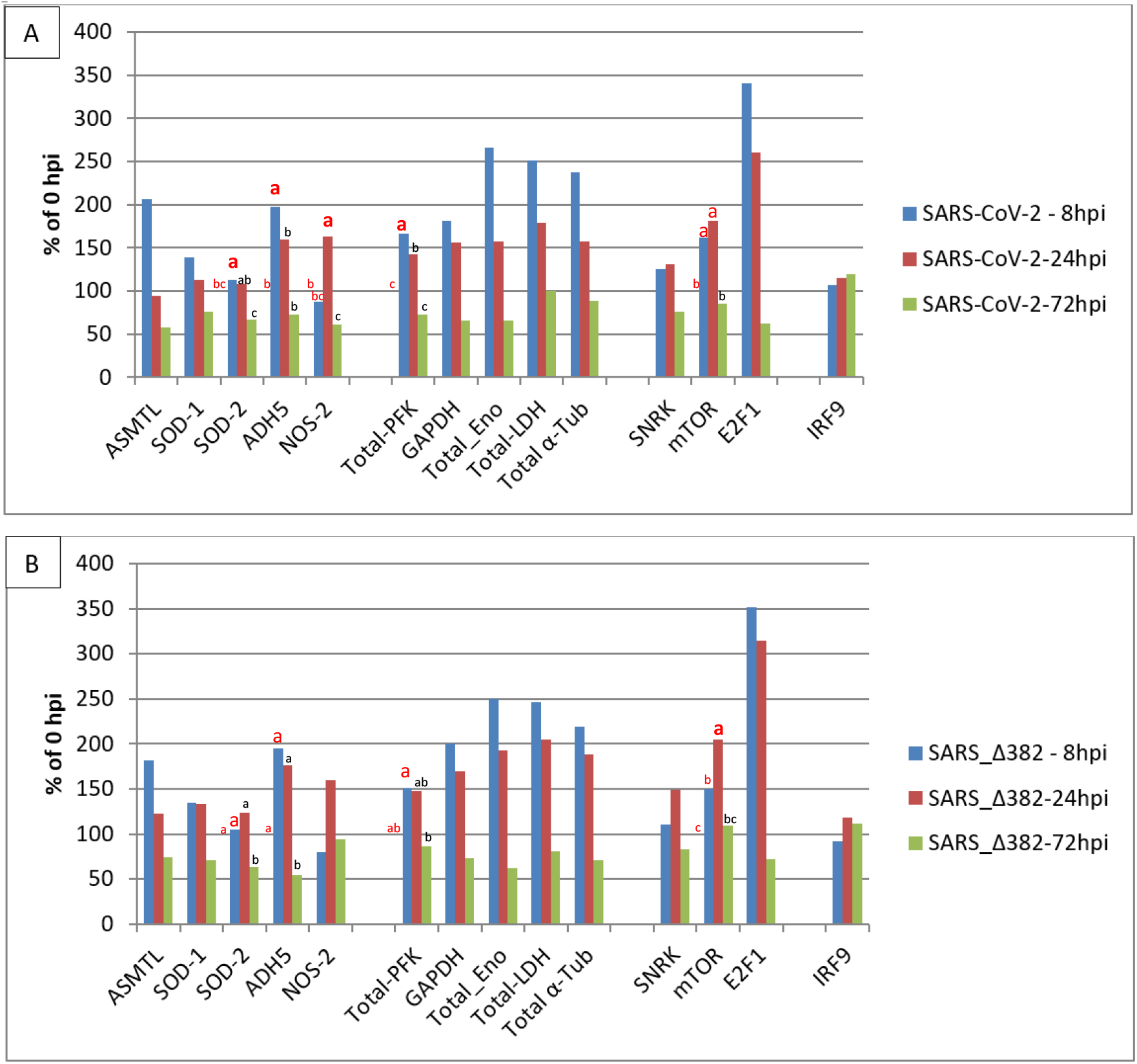
Transcript accumulation of selected *ReprogVirus* marker genes in human nasal epithelial cells infected with two SARS-CoV-2 variants at 8 hours post infection (hpi), 24 hpi and 72 hpi. Transcript levels are averages from three cell origins (donators/cell cultures) given in % of 0 hpi. A – SARS-CoV-2 (virus originally discovered); B – SARS-CoV-2 Δ382. 2-way ANOVA analysis identified differential transcript level changes along early times between the original virus and the mutant (marked by enlarged, fat letters). Different letters indicate significant differences between net RPKM for α = 0.05. Letters on the 100% horizontal line correspond to 0 hpi. The result of 2-way ANOVA analysis for all genes can be consulted in **supplementary Figure S1**).

Transcript profile level changes for SARS-CoV-2 (original virus) infection are given in Fig.1A and demonstrate similar increases for ASMTL and ADH5 (207% and 197%) at 8 hpi and indicate unbalanced ROS/RNS at 24 hpi (94% ASMTL and 159% ADH5), which is supported by near-basal SOD1 and SOD2 transcript levels (112% and 108%) in relation to an increased transcript level of NOS2 (163%). At 72 hpi, transcript levels of all ROS/RNS related genes were simultaneously reduced to below basal values observed at 0 hpi (57% ASMTL, 76% SOD1, 68% SOD2, 73% ADH5 and 61% NOS2). Transcription of glycolysis-related enzymes was rapidly increased at 8 hpi with highest transcript accumulation for enolase (267%). This high increase at 8 hpi was associated to a similar increase in LDH transcription (251%) linked to highly enhanced transcript levels also for α-tubulin (238%). SARS-CoV-2 Δ382 (Fig. 1B) shows a very similar profile for ROS/RNS signaling and metabolism-and structural cell organization-related traits. They also display almost identical signaling for a change in cell cycle regulation and the lack of strong stimulation of the classical immune system represented by the missing of IRF9 transcript accumulation level changes. Both genotypes demonstrate high increase in E2F1 transcript stimulation at 8 hpi, which signals rapid cell cycle activation. E2F1 transcript levels decreased slightly at 24 hpi, but fall rapidly below the basal values observed at 0 hpi to 62% respectively 72% (at 72 hpi). These observations together with increased transcript levels of SNRK and mTOR at 8 hpi (126% and 162%) and 24 hpi (131% and 182%) for the original SARS-CoV-2, and for SARS-CoV-2 Δ382 at 8 hpi (111% and 150%) and at 24 hpi (149% and 205%) point for both virus variants to early energy depletion and rapidly driven cell cycle progression plus cell cycle arrest at 72 hpi. At 72 hpi, no energy depletion is signaled anymore for both virus variants (SNRK: 76%, 83%). However, SARS-CoV-2 Δ382 shows at 72 hpi a slower decrease of mTOR transcript accumulation (109% vs 86% for original virus), and this together with the slower decrease also for E2F1 transcription (72% vs. 62% for original virus) might indicate slightly delayed cell cycle progression and arrest for the mutant. These last observations link to the postponed increase in transcript levels of LDH (at 8 hpi 247% mutant vs 251% original virus and at 24 hpi 205% mutant vs 179% original virus) and α-Tub (at 8 hpi 220% mutant vs 238% original virus and at 24 hpi 189% mutant vs 157% original virus). Further, considering delayed cell cycle progression for SARS-CoV-2 Δ382 is supported by 2-ways ANOVA analysis. This analysis identified significant transcript level increases of SOD2, ADH5 and PFK early at 8 hpi and highlighted a significant increase for NOS2 from 8 hpi to 24 hpi only for the original virus. On the other hand, it demonstrated a significant increase for mTOR from 8 hpi to 24 hpi only for the mutant. However, the effect of such differences between original virus and the mutant did not substantially influence the initiation kinetics of IRF9 transcription. Transcript level increase for IRF9 was marginally seen at 72 hpi for both virus variants (>110%, both non-significant) and only a slightly different extent is indicated between original virus and mutant (120% vs 111%). The infection trials had been performed in unsynchronized nasal cell cultures (48, 58). Thus, it cannot be excluded that the observed differences in transcript levels between both virus variants may be due to experiment-dependent, differentially non-synchronized cell cycle phases.

**Figure 2** shows transcript accumulation in RPKM for both virus variants from nasal epithelial cells separated for the three cell origins that resulted in the averages shown in Figure 1. Additionally, this figure integrates transcript accumulation of virus helicase and virus RNA-dependent RNA-polymerase (RdRp) as markers for virus proliferation. In **Figure 2A**, it can be seen that cells from origin 1 demonstrate lower transcript levels for enolase and LDH at 24 hpi. This was observed after infection by both SARS-CoV-2 variants. Also for both virus variants, virus helicase and RdRp indicate the start of virus replication at around 24 hpi though to a slightly lower degree for the SARS-CoV-2 Δ382. In **Figure 2B**, transcript accumulation of virus helicase and virus RdRp point to a burst of virus replication at 72 hpi. The level of virus transcripts is lower for the mutant than for the original SARS-CoV-2 virus. However, cells that originated from donator 1 that showed lower transcript accumulation for enolase and LDH at 24 hpi in comparison to cells from donators 2 and 3 in response to both virus variants did not indicate a burst of virus replication at 72 hpi, but only a slow increase from 24 hpi. Nevertheless, cells of donator 1 also demonstrated lower virus RdRp transcript levels at 72 hpi (**Fig. 3**). This differential performance of cells from donator 1 connects to higher transcription levels of SNRK at 72 hpi in cells from donator 1 after infection with both, the original SARS-CoV-2 and SARS-CoV-2 Δ382 (**Fig. 2 and 3**). It signals higher depletion of energy for donator 1 cells at 72 hpi and confirms critical energy-dependency of SARS-CoV-2 replication for both variants. However, the higher values for enolase and LDH observed at 24 hpi for cells infected by the mutant, the differential development of SNRK over time and higher values at 72 hpi observed for mTOR from donator 1 in the mutant (**Fig. 3**) argues for prolonged cell cycle progression respectively a delay in cell cycle arrest related to both, the origin of cells from donator 1 and infection by the mutant. Nevertheless, since longer observation times and technical repetitions in synchronized cultures are missing, it cannot be concluded whether individual origin or differentially unsynchronized cell cultures caused these differences in the emergence of a burst in virus replication.

**Figure 2:**
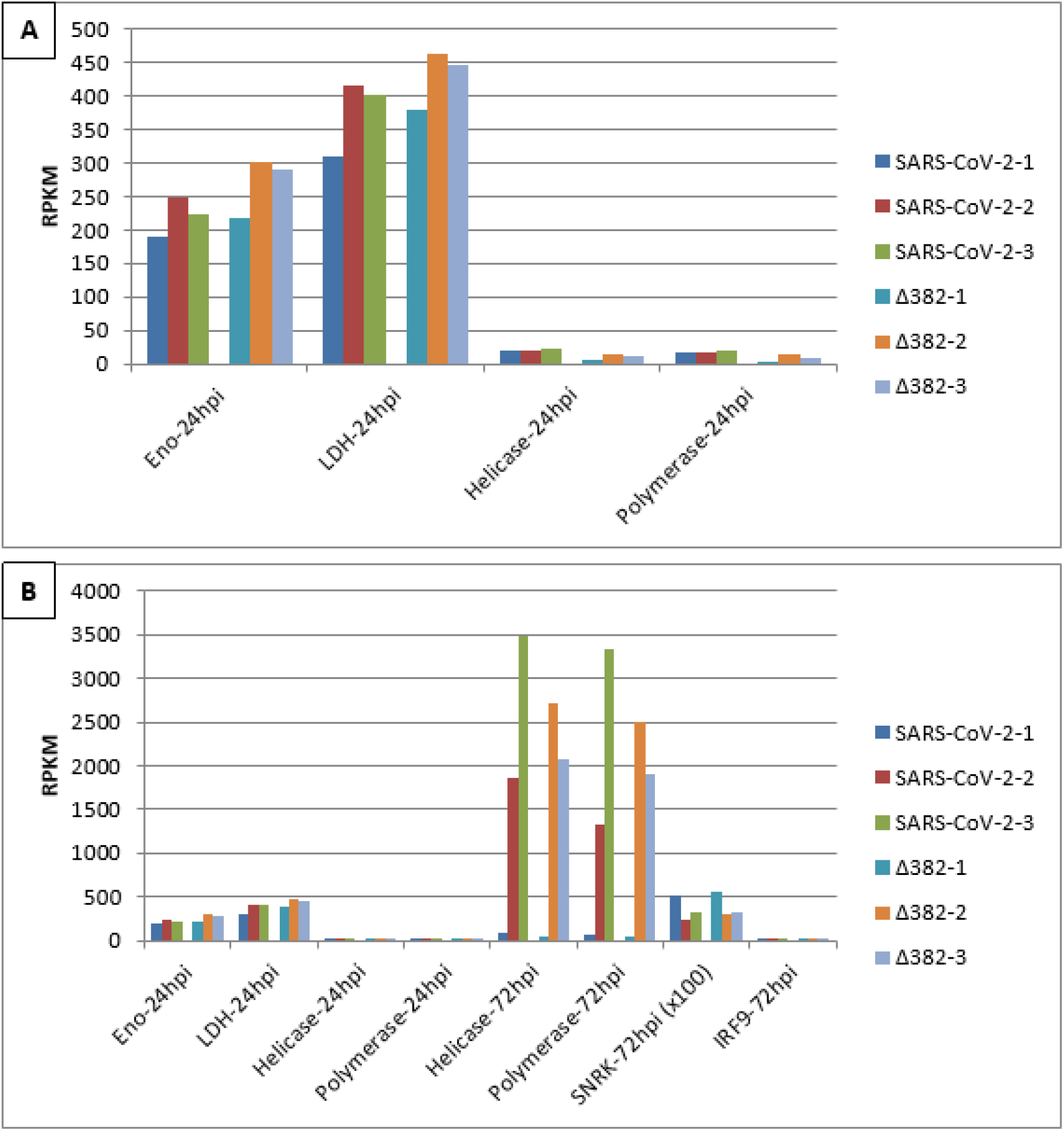
Burst of SARS-CoV-2 virus proliferation at 72 hpi in nasal epithelial cells from three origins infected by two SARS-CoV-2 variants indicates an energy-dependent link to aerobic glycolysis and fermentation at 24 hpi.

**Figure 3:**
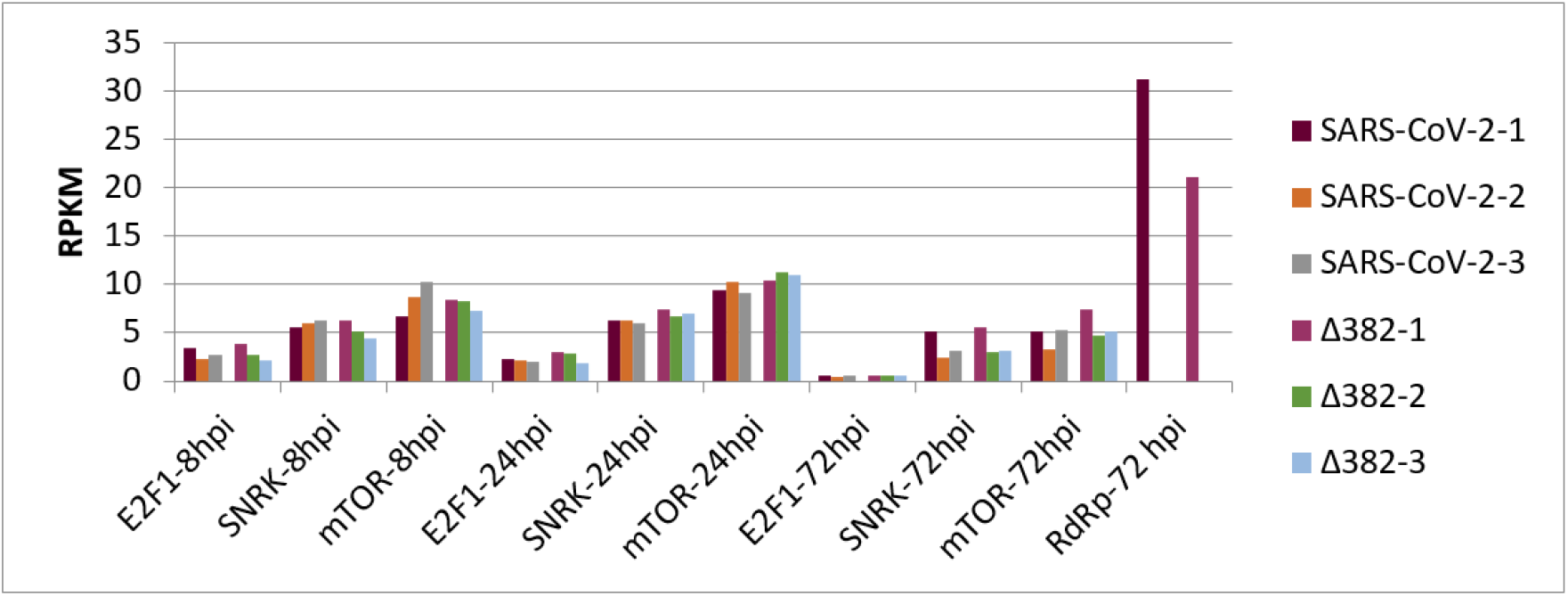
Transcript accumulation of E2F1, SNRK, and mTOR in human nasal epithelial cells infected by two SARS-CoV-2 variants at 8 hpi, 24 hpi and 72 hpi and transcript levels of RdRp (x 0,5) at 72 hpi for cell origin 1

## 4. Discussion

This is the first time that relevance of the marker system ‘CoV-MAC-TED’, which is based on a relatively small set of key genes, was validated in primary human target nose cells for respiratory SARS-CoV-2 infections. It is also the first time that this approach was tested for two SARS-CoV-2 variants, which associate to differences in clinical effects (48, 49).

Our analyses have been performed in public RNA sequencing data provided from experiments with human NECs described in Tan et al. (58) and Gamage et al. (48). Thus, results and conclusions of these authors helped us additionally to validate relevance our key marker candidate approach: Transcriptome accumulation of the selected set of ‘ReprogVirus’ genes showed similar performance of ‘CoV-MAC-TED’ in human NECs infected by the originally discovered SARS-CoV-2 virus compared to infection by SARS-CoV-2 Δ382. In this aspect, our results are in conformity with the conclusions drawn by Gamage et al. (48). In addition, our observation that both virus variants induced cell cycle activator E2F1 to highest transcript levels at 8 hpi among all tested genes and times is in good agreement with results of Gamage et al. (48). These authors highlighted enhanced numbers of transcription factor E2F targets during early virus infections and a decrease during time combined with high activity related to cell cycle checkpoint G2M.

However, applying CoV-MAC-TED indicated also differences between both SARS-CoV-2 variants. The complex marker components revealed virus-induced ROS/RNS de-balancing, differential glycolysis, fermentation and cell cycle regulation that pointed to delayed cell response and cell cycle arrest for the mutant, which connected to delayed virus propagation. Interestingly, a delay in cell cycle arrest was also indicated for one of three cell origins (cells from donator 1), which pointed to the relevance of lower levels of glycolysis and aerobic fermentation for postponing virus replication (Figure 2).

In the same RNAseq data that we used for the presented research, the providing authors Gamage et al. (48) had intensively studied the complex immune response of SARS-CoV-2 infected NECs compared to cells infected by influenza H3N2. Influenza infection showed an earlier burst of virus replication at 48 hpi related to a pronounced early initiated immune response. This gave us the opportunity to further validate, advance and standardize our approach in a parallel study with a focus on influenza H3N2 infected NECs (59, preprint). In this way, we could validate our choice of IRF9 (12) as an appropriate general marker for the classical immune system. The usefulness of CoV-MAC-TED to identify similar and differential early cell responses was strengthened. Both virus types showed early unbalanced ROS/RNS and temporarily increased aerobic fermentation linked to α-tubulin-marked cell restructuration. However, CoV-MAC-TED indicated the absence of initial cell cycle progression for influenza A H3N2 infections that connected to rapid energy-dependent IRF9-marked immunization and this contrasted our present findings during infections with both SARS-CoV-2 variants.

In **Figure 4**, we present a simplified scheme from the results here obtained in human NECs infected by SARS-CoV-2 for the five main components of CoV-MAC-TED (1) ROS/RNS balance shift, represented by ASMTL, SOD1, SOD2, ADH5, and NOS2, (2) glycolysis and fermentation, represented by enolase and LDH, (3) structural reorganization and cell cycle progression and arrest, represented by E2F1, mTOR and α-tubulin, (4) energy status signaling, represented by SNRK and mTOR, and (5) initiation of the classical immune system, represented by IRF9. For our conclusions and hypothetical inferences, we consider the dynamic interplay between virus infection, ROS/RNS signaling, carbohydrate stress metabolism, aerobic fermentation and cyt respiration based on knowledge related to our recently published insights (11, 26) and state-of-the-art knowledge on SARS-CoV-2 infection and general stress biology. Thus, we expect that the indicated increase of oxidative stress signaled at 8 hpi by ASMTL and ADH5 upon SARS-CoV-2 infection associates with rapidly increased sucrose/glucose cell levels that stimulate the Cyt pathway via enhanced glycolysis, pyruvate production and increased TCA cycling in a way that the respiration chain gets overloaded by electrons. Consequently, ROS and RNS concentrations might increase. On the other hand, the Cyt pathway will temporarily be restricted due to rapidly consumed oxygen. In turn, aerobic lactic acid fermentation will be activated. Depending on stress level and the amount of sugar and duration of the situation of high-sugar level, anaerobic glycolysis can reach high turnover during cell reprogramming and a level of ATP production corresponding to the Warburg effect. Warburg effects are increasingly recognized as being part of normal physiology (60, 61) that enables host cells to rapidly mobilize energy for host maintenance and stress escape. Results of Bharadwaj et al. (26) suggests that microbiota can help stress alleviation through forming a sink for sucrose that supports their own growth, and, depending on their nature, this could support symbiotic or parasitic performance. Viruses are parasitic structures. These structures are supposed to profit from the mobilized high energy favored by the thermodynamic conditions of their replication. However, in case of SARS-CoV-2 our results indicate that during the first hours post infection (8 hpi to 24 hpi) rapidly available energy is consumed first for cell cycle progression (E2F1, mTOR and α-tubulin signaling) and only from around 24 hpi energy is increasingly used for virus replication (Figure 2). This might signal during early hours after infection stress (observed at 8 hpi by highly increased ASMTL and ADH5 transcription) relieve to the host cell (low oxidative stress indicated by ASMTL, SOD1 and SOD2), lower sugar concentration (less glycolysis combined with lower degree of fermentation observed from 24 hpi) and, thus, enable normalization of TCA cycling and decrease of energy depletion as it is SNRK-signaled after 24 hpi at 72 hpi before the expected burst of the classical immune system. However, the change for ROS/RNS in favor of NO production at 24 hpi, when virus replication starts, points to a ‘non-normal’ unbalanced situation.

**Figure 4:**
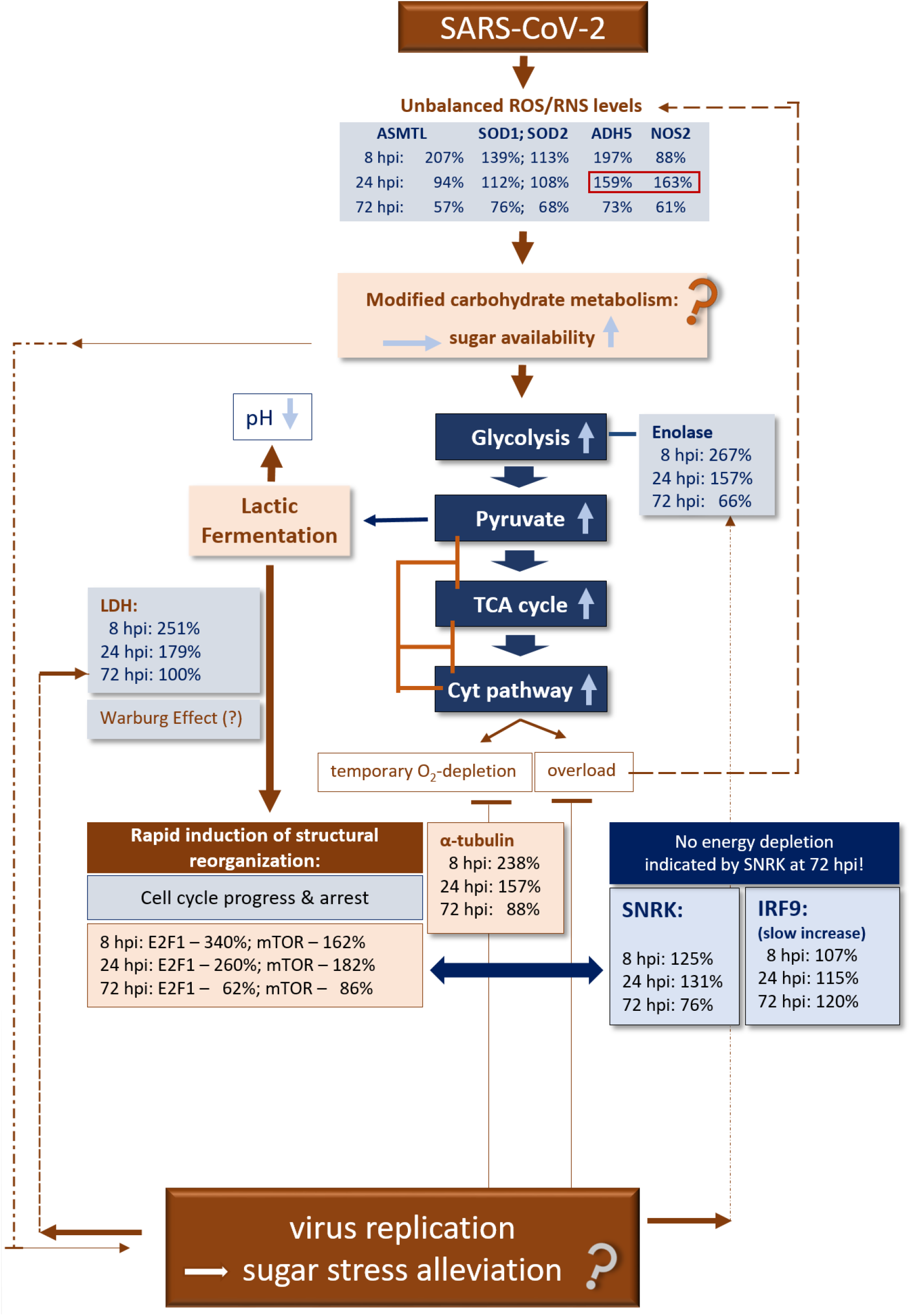
Validating CoV-MAC-TED as crucial trait for early SARS-CoV-2-induced reprogramming in human nasal epithelial cells – A simplified scheme for hypothetical metabolic principles and validation of markers.

## Conclusion

The high relevance of metabolism-driven, early cell re-organization that we observed by studying CoV-MAC-TED in infected primary target cells for two respiratory viruses, SARS-CoV-2 and influenza A H3N2, stimulate re-thinking of current understanding of the immunological system and its determinants. Future anti-viral effective therapeutic strategies should, to our view, explore more sensitive diagnostic virus tools for rapid virus identification combined with effective individual treatment through early targeting CoV-MAC-TED components. Lower transcript levels observed for enolase and LDH observed at 24 hpi for cells/cell cultures from one origin that was linked to delayed virus replication might point to a strategic target for combating early initiation of SARS-CoV-2 replication by blocking the Warburg effect and its link to cell cycle progression.

*In vitro*-cultured human nasal epithelial cells proved to be appropriate for standardized transcriptome data-collection in the ‘ReprogVirus Platform’. Thus, this cell system is highly promising to advance integrative data analyses by help of Artificial Intelligence methodologies for designing anti-SARS-CoV-2/anti-viral strategies.

## Author Contribution

JHC & BA-S conceived the basic idea and plan of the study. SA performed BLAST and helped JHC in transcriptome analyses and manuscript preparation. CN performed statistical analyses. BA-S reviewed data analyses, interpretation, wrote the manuscript and coordinated revision of the manuscript. All co-authors commented and approved the final manuscript.

## Funding

JHC is grateful to CNPq for the Researcher fellowship (CNPq grant309795/2017-6). SA is grateful to CAPES for the Doctoral fellowship.

## Acknowledgements

We would like to express special appreciation to our colleagues from Singapore, who established NECs from individual cell origins and provided public RNAseq data (Tan et al., 2018 and Gamage et al., 2019), which was very helpful to effectively advance our research approach and line of strategic goals. The authors would like to thank Isabel Velada from the University of Évora, Portugal, for help in figure improvement. Furthermore, BAS wants to acknowledge the availability of Arvind Achra, Department of Microbiology, Atal Bihari Vajpayee Institute of Medical Sciences & Dr Ram Manohar Lohia Hospital, New Delhi, India, to discuss feasibility of future strategies to combat SARS-CoV-2 infections on the basis of his work experience in a laboratory for diagnosis.

## Conflicts of Interest

The authors declare that there is no conflict of interest.

## Notes

### Competing Interest Statement

The authors have declared no competing interest.

### Summary of Updates

This version of the manuscript was revised during the review process provided by the journal Vaccines. Also, we integrated an additional scientist as co-author in order to help improving the manuscript by further statistical analysis.

